# Identifying SARS-CoV-2 Antiviral Compounds by Screening for Small Molecule Inhibitors of Nsp15 Endoribonuclease

**DOI:** 10.1101/2021.04.07.438811

**Authors:** Berta Canal, Ryo Fujisawa, Allison W. McClure, Tom Deegan, Mary Wu, Rachel Ulferts, Florian Weissmann, Lucy S. Drury, Agustina P. Bertolin, Jingkun Zeng, Rupert Beale, Michael Howell, Karim Labib, John F.X Diffley

**Affiliations:** Chromosome Replication Laboratory, The Francis Crick Institute, 1 Midland Road, London, NW1 1AT; High Throughput Screening, The Francis Crick Institute, 1 Midland Road, London, NW1 1AT; Cell Biology of Infection Laboratory, the Francis Crick Institute, 1 Midland Road, London, NW1 1AT; The MRC Protein Phosphorylation and Ubiquitylation Unit, School of Life Sciences, University of Dundee, Dundee DD1 5EH, UK

## Abstract

SARS-CoV-2 is responsible for COVID-19, a human disease that has caused over 2 million deaths, stretched health systems to near-breaking point and endangered the economies of countries and families around the world. Antiviral treatments to combat COVID-19 are currently lacking. Remdesivir, the only antiviral drug approved for the treatment of COVID-19, can affect disease severity, but better treatments are needed. SARS-CoV-2 encodes 16 non-structural proteins (nsp) that possess different enzymatic activities with important roles in viral genome replication, transcription and host immune evasion. One key aspect of host immune evasion is performed by the uridine-directed endoribonuclease activity of nsp15. Here we describe the expression and purification of nsp15 recombinant protein. We have developed biochemical assays to follow its activity, and we have found evidence for allosteric behaviour. We screened a custom chemical library of over 5000 compounds to identify nsp15 endoribonuclease inhibitors, and we identified and validated NSC95397 as an inhibitor of nsp15 endoribonuclease *in vitro*. Although NSC95397 did not inhibit SARS-CoV-2 growth in VERO E6 cells, further studies will be required to determine the effect of nsp15 inhibition on host immune evasion.

## Introduction

SARS-CoV-2 (severe-acute-respiratory-syndrome coronavirus-2) is the coronavirus that causes the human coronavirus disease (COVID-19) [1, 2]. SARS-CoV-1 and MERS-CoV caused severe diseases in 2003 and 2012, respectively, and, although their consequences in terms of worldwide loss of human life cannot be compared to the devastating health and economic crisis caused by COVID-19 pandemic, the emergence of SARS-CoV-2 indicates a pattern of recurrent coronavirus diseases with relevance for human health [3, 4]. Some features of SARS-CoV-2 have been pinpointed as key to explain the catastrophic consequences of COVID-19; an easy air-transmissibility, a large percentage of asymptomatic infections, and the ability of the virus to infect multiple cell types and organs and deregulate the immune system [5–9].

In a major international effort, multiple vaccines are being developed and several are already being administered. It is still uncertain whether vaccines will fully prevent future infections or lead to milder COVID-19 infections [10]. Most vaccines are being developed against the spike protein of the virus, a structural protein that evolves rapidly and, therefore, might not be a long-term solution to COVID-19 and, moreover, is unlikely to provide a pan-coronavirus solution [11, 12]. There is currently a lack of antiviral treatments that efficiently combat coronavirus infections. Remdesivir, a nucleotide analogue chain terminator proven to reduce viral growth, is the only antiviral approved for the treatment of COVID-19 [13]. However, a WHO-funded clinical trial recently failed to demonstrate a reduction in deaths amongst patients treated with remdesivir [14]. Therefore, the further development of efficient antiviral treatments to combat diverse coronavirus infections will remain important for health systems worldwide to face present and future coronavirus pandemics.

SARS-CoV-2 is a (+) single-strand RNA virus that belongs to the family *Coronaviridae* of the order *Nidovirales*, which includes viruses with the largest known RNA genomes [15–17]. The first two-thirds of the genome encode two open-reading frames, ORF1a and ORF1b, that are directly translated from the viral genome producing two large polyproteins (pp1a and pp1ab) that contain the 16 non-structural proteins (nsps) of the virus. These polyproteins are cleaved by the proteolytic activities of nsp3 and nsp5, which therefore modulate the activity of the rest of the enzymes and promote the formation of active protein complexes including the RNA-dependent RNA polymerase (nsp12/nsp7/nsp8), the RNA helicase (nsp13), the RNA exoribonuclease (nsp14/nsp10), the RNA endoribonuclease (nsp15) and the RNA Cap methyltransferases (nsp14 and nsp16/nsp10), which together support proliferation of the virus. Therefore, coronaviruses contain a wide variety of enzymes that are potential targets for the development of novel antivirals, by comparison with many other viruses.

nsp15 contains a C-terminal ‘EndoU’ domain (*endo*ribonuclease *u*ridylate-specific) that is able to cleave the 3’ end of pyrimidines, preferentially uridylates, in the context of single and double stranded RNA molecules [18–21]. Purified nsp15 can exist as an inactive monomer or trimer, or an active hexamer [22, 23]. Oligomerisation relies on N-terminal residues of nsp15 and activity is dependent on the presence of divalent metal ions, with a preference for Mn^2+^ [19, 23, 24]. The EndoU activity of nsp15 is dispensable for replication of viral RNA genome [20, 25]. Instead, nsp15 EndoU activity appears to mediate the evasion of host recognition of viral dsRNA, and growth of EndoU-deficient viruses is severely affected in mouse models and primary immune cells such as macrophages [21, 26, 27]. Therefore, nsp15 EndoU inhibitors might be useful to potentiate the host immune response to SARS-CoV-2.

In March 2020, we initiated a large project to identify inhibitors of multiple SARS-CoV-2 enzymes from a custom chemical library containing over 5000 compounds. Here we describe the results of a screen for nsp15 endoribonuclease inhibitors.

## Results

### Purified SARS-CoV-2 nsp15 has uridylate-specific endoribonuclease activity in vitro

We expressed and purified a series of N-terminally tagged versions of nsp15 from bacteria and insect cells (Figure 1A). Nsp15 eluted as a high molecular weight complex during size-exclusion chromatography, consistent with nsp15 existing as a hexamer [22, 23] (Supplementary Figure S1A). Removal of the affinity tags with proteases was generally inefficient, consistent with a recent structural study that showed that the N-terminus of nsp15 is not on the surface of the nsp15 hexamer [28]. After partial digestion of 14His-SUMO-nsp15 with Ulp1 SUMO protease, untagged nsp15 eluted much later during size-exclusion chromatography, suggesting the untagged protein was primarily present in a smaller complex or as a monomer (Supplementary Figure S1A and S1B).

**Figure 1.**
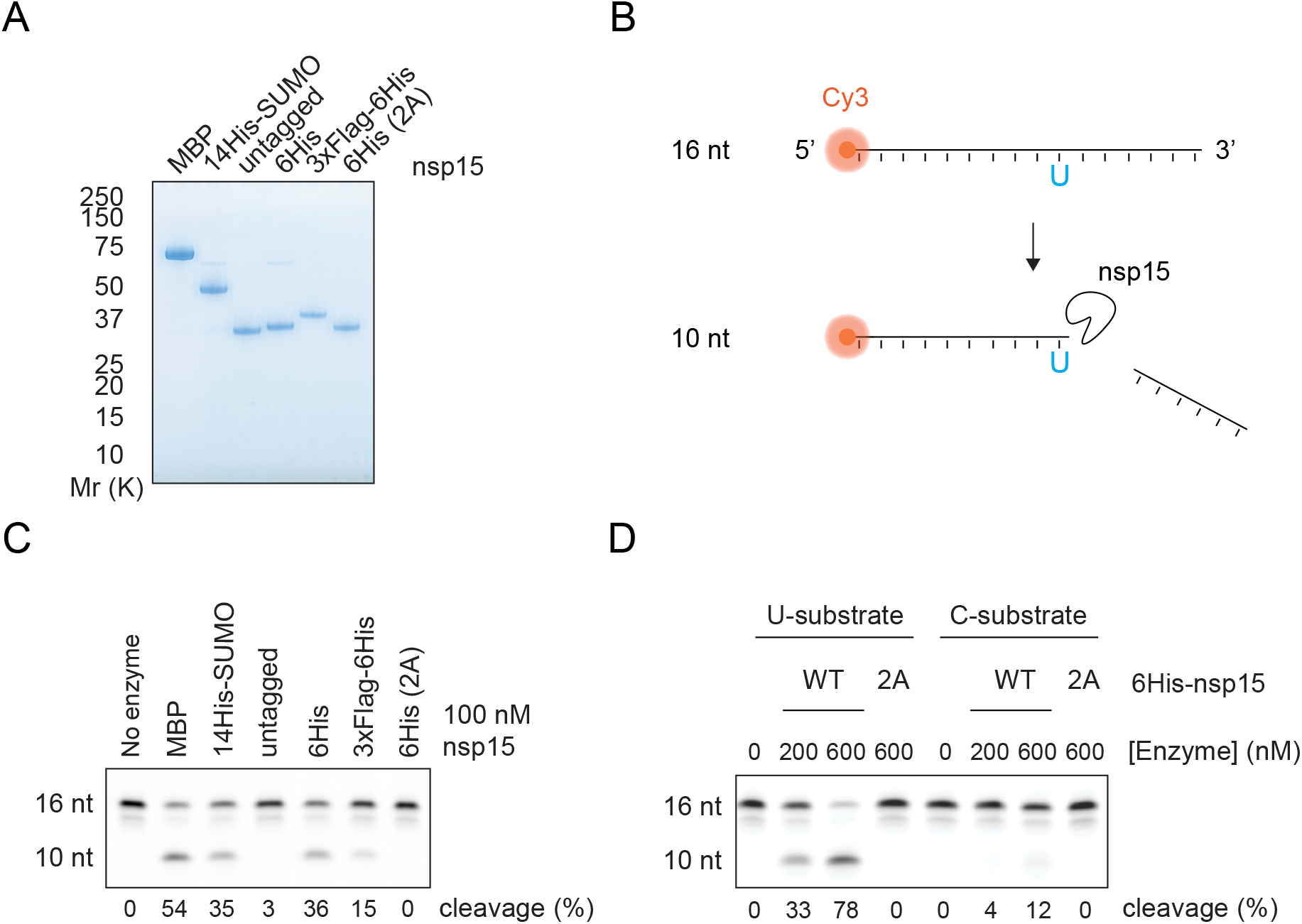
Purification of active SARS-CoV-2 nsp15 endoribonuclease. **A**. A range of affinity tags were used to purify nsp15. Purified proteins were visualised by SDS-PAGE and Coomassie Blue staining. ‘untagged’ nsp15, which was generated by partial proteolytic cleavage of 14His-SUMO nsp15 with Ulp1 SUMO protease, eluted much later during size-exclusion chromatography, consistent with the untagged nsp15 existing as a smaller complex or monomer (see Supplementary Figure S1). 6His (2A) protein corresponds to the H234A H249A mutant of nsp15. **B**. Scheme of the gel-based endoribonuclease assay for nsp15. A 16 nt ssRNA containing a 5’ Cy3 fluorophore (16 nt U substrate) is cleaved at the only uridine (U) to yield a 10 nt product. A version of this substrate with a cytosine instead of uridine is used to test specificity of nsp15 cleavage in D (16 nt C substrate). **C**. Cleavage of the 16 nt U substrate by the different purified nsp15 proteins shown in A. 100 nM of nsp15 and 1 μM of substrate were incubated for 30 min at 30°C and then resolved in a denaturing TBE-urea polyacrylamide gel. Quantification of the percent product in the cleaved 10 nt band is given at the bottom of the gel. **D**. The U or C version of the 16 nt substrate (1 μM) were incubated with different concentrations of wildtype (WT) or 2A-endoribonuclease mutant 6His-nsp15, and endonuclease reactions were performed as in C.

To evaluate nsp15 endoribonuclease activity, we used a 16 nt single stranded RNA substrate labelled with a Cy3 fluorophore, which contained a single uridine near the centre (Figure 1B). N-terminal MBP-, 14His-SUMO-, 6His- and 3xFLAG-6His- tagged versions of nsp15 were able to cleave this substrate, whereas the nsp15-2A mutant in which two key Histidine residues (H234A H249A) were replaced with alanine, did not [29] (Figure 1C). Notably, untagged nsp15 generated from the digestion of 14His-SUMO-nsp15, was inactive as an endoribonuclease consistent with the idea that only the hexameric form of nsp15 is active [22, 23, 30]. We selected 6His-nsp15 for use in subsequent screening and validation assays. Whilst this enzyme was active on the uridylate-containing substrate, it had no detectable activity on a substrate in which the single uridine was substituted for cytidine (Figure 1D). Taken together, these results show that the purified 6His-nsp15 possesses uridylate-specific endoribonuclease activity, consistent with previous data [22, 23, 30].

### Assay to study kinetic SARS-CoV-2 nsp15 endoribonuclease activity in solution

To perform a high-throughput screen for inhibitors of nsp15, we first developed an assay to quantify nsp15 endoribonuclease activity in solution without the need to visualise products in a gel. We used a short (6 nt) single stranded RNA oligonucleotide containing a single uridine and a 5’ Cy5 that was quenched by proximity to a 3’ BBQ-650 quencher. This substrate only exhibits fluorescence upon cleavage by nsp15, when the Cy5 is no longer in proximity to the quencher (Figure 2A). We found that nsp15 was able to cleave the 16 nt (Figure 2B, top) and 6 nt (Figure 2B, bottom) substrates at similar enzyme concentrations. As expected, fluorescent signal from the 3 nt cleavage product was only detectable upon substrate cleavage. The uridylate specificity of nsp15 was confirmed again by its lack of activity on a 6 nt substrate containing cytidine in place of uridine (Figure 2C).

**Figure 2.**
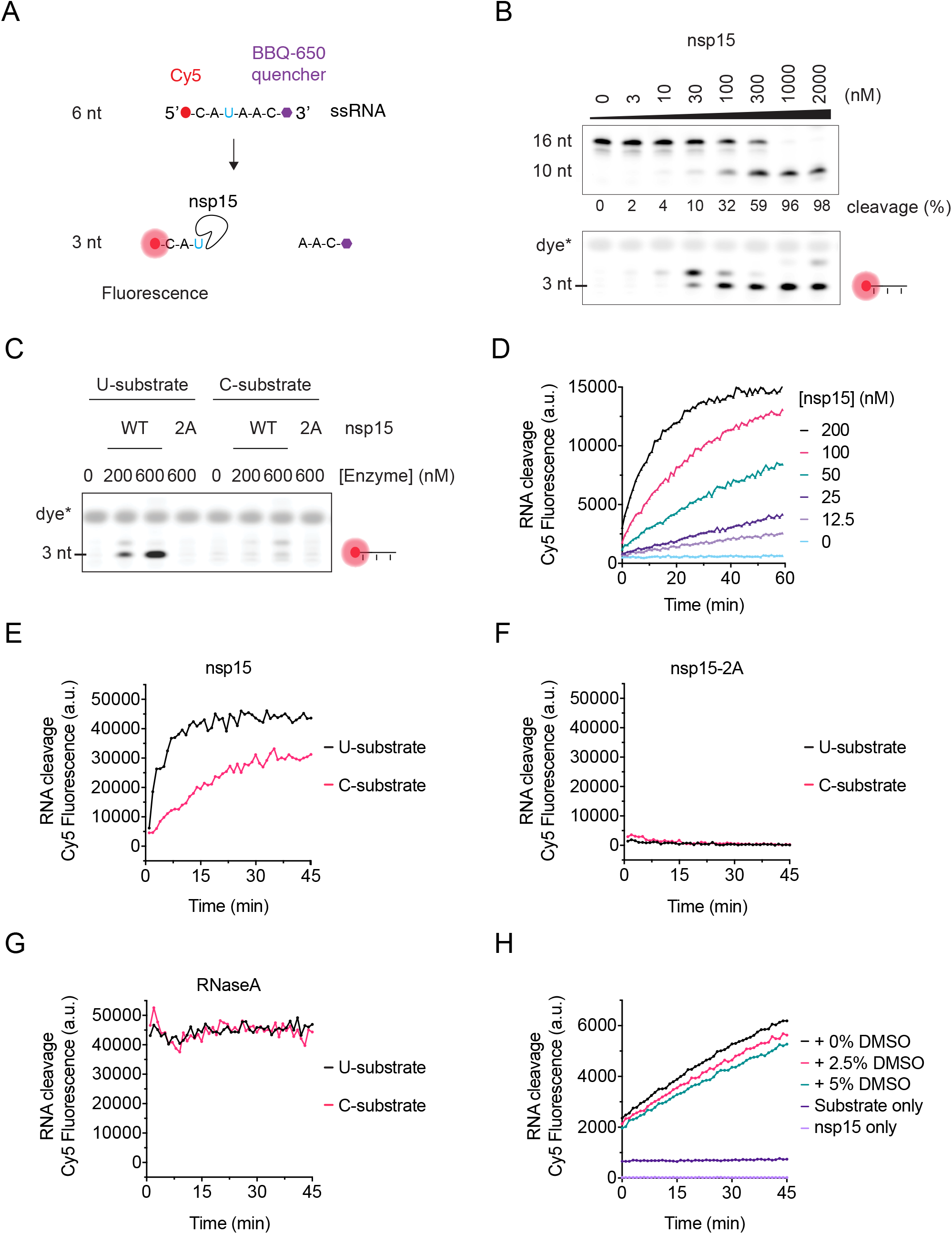
Development of a fluorescent biochemical nsp15 endoribonuclease assay. **A**. A 6 nt ssRNA containing a 5’ Cy5 fluorophore quenched by a 3’ BBQ-650 quencher (6 nt U substrate) was used to assess the activity of nsp15 in gels-based assays and in solution. Upon addition of the nsp15 endoribonuclease, cleavage of the oligo at its only uridine (U) releases the quencher, leading to Cy5 fluorescence. A version of this substrate with a cytidine in place of the uridine (6 nt C substrate) is used to test the uridine specificity of nsp15 cleavage. **B**. Titration of nsp15 enzyme in the presence of 1 μM of the 16 nt U substrate (upper gel) or 1 μM of the 6 nt U substrate (lower gel) showing activity on both substrates at similar enzyme concentrations. Reactions were performed for 30 min at 30°C and resolved in a denaturing TBE-urea denaturing polyacrylamide gel. Fluorescence of the 6 nt substrate was only observed upon cleavage by nsp15. Appearance of a fluorescent band above the 3 nt suggests that nsp15 can cleave the 6 nt substrate at more than one residue. **C**. The “U” and “C” version of the 6 nt substrate (1 μM) were incubated with different concentrations of wildtype (WT) or 2A-endoribonuclease mutant 6His-nsp15 enzyme and endonuclease reactions were performed as in B. **D**. Titration of nsp15 enzyme (0 – 200 nM nsp15) in the presence of 200 nM of the 6 nt U substrate performed at RT and fluorescence quantified in a Spark Multimode microplate reader (Tecan) every min for a total of 60 min. **E**. Cleavage of the “U” and “C” versions of the 6 nt substrate (1 μM) in the presence of 500 nM nsp15. Reactions were performed at RT and fluorescence was measured in an infinite M1000 Pro reader (Tecan) every min for 45 min. **F.** Same reactions as in E performed with 500 nM of nsp15-2A mutant. **G.** Same reactions as in E performed with 0.1 mg/ml RNase A. **H**. Cleavage of the 6 nt U substrate (500 nM) in the presence of 75 nM nsp15 and 0, 2.5% or 5% DMSO, and fluorescence measured in a Spark Multimode microplate reader (Tecan) every min for 45 min. Fluorescence of reactions containing substrate-only and enzyme-only controls is also shown.

Prior to performing the screen to identify inhibitors of SARS-CoV-2 nsp15, we adapted the in-solution fluorescent biochemical assay to a multi-well plate format. We first performed a titration of nsp15 and were able to detect Cy5 fluorescence on a plate reader in a time and enzyme concentration-dependent manner (Figure 2D). As seen in gel-based assays, nsp15 exhibited reduced endonuclease activity toward the C substrate when using the plate reader (Figure 2E) and, accordingly, the nsp15-2A catalytically inactive mutant did not produce fluorescent signal on either substrate (Figure 2F). In parallel, the nuclease RNase A showed efficient cleavage of both substrates (Figure 2G). As the compounds in the chemical library to be used for the screen were stored in DMSO, we also tested nsp15 activity in the presence of DMSO and found DMSO at concentrations up to 5% had only a slight inhibitory effect on enzyme activity (Figure 2H).

### Fluorescent biochemical kinetic screen for SARS-CoV-2 nsp15 inhibitors

We next used this assay to determine the effect of substrate concentration on initial reaction velocities. We found that, in contrast to all of the other enzymes we have examined in this series of papers (see accompanying manuscripts), reaction rates did not plateau at higher substrate concentrations but instead continued to increase non-linearly, indicating that nsp15 endoribonuclease behaves as an allosteric enzyme towards the 6 nt U substrate, with a K_half_ of 2140 nM (Figure 3A and Supplementary Figure S2A).

**Figure 3.**
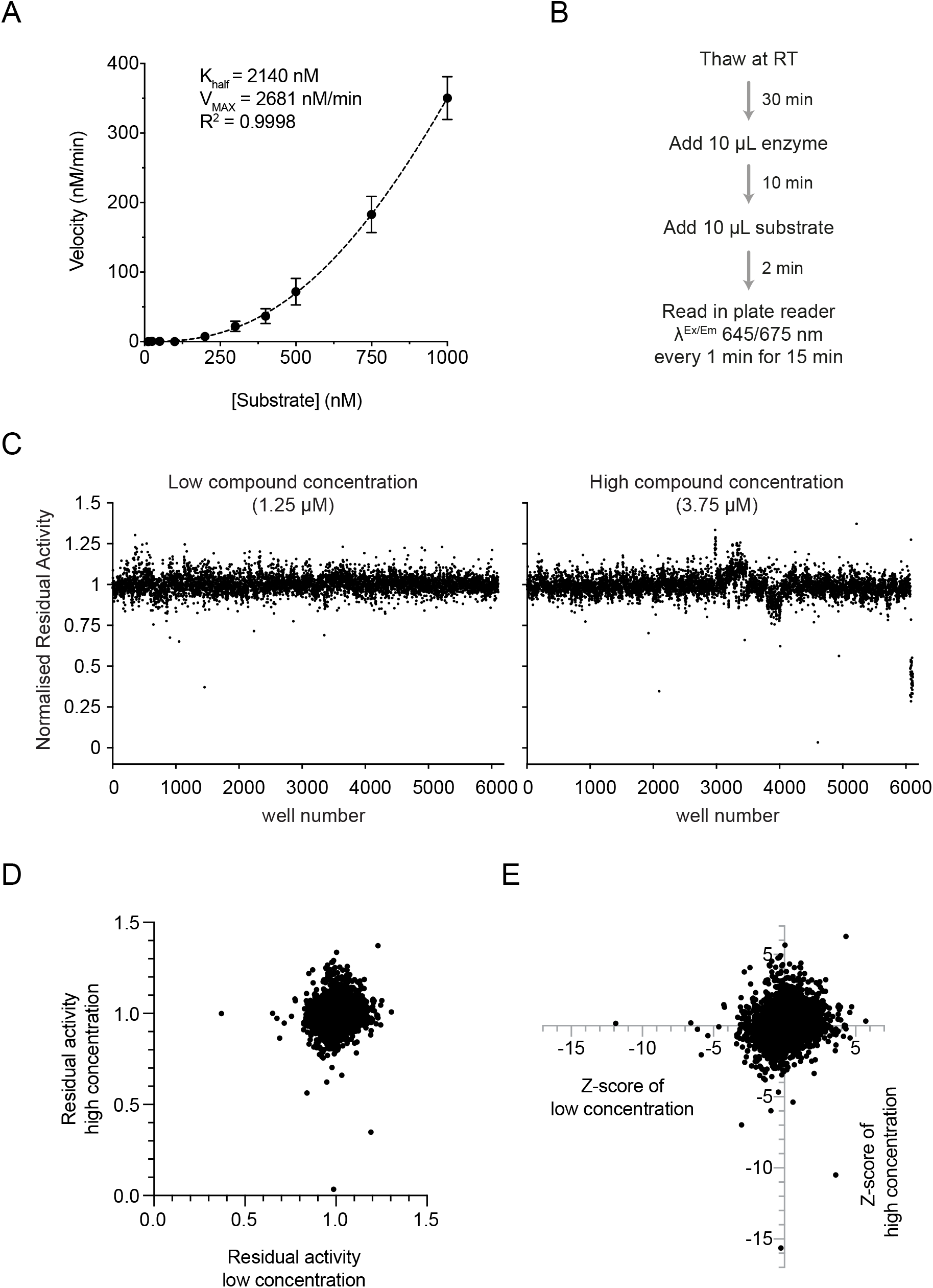
SARS-CoV-2 endoribonuclease nsp15 inhibitor screen design and results. **A.** Non-linear fit to an Allosteric sigmoidal equation of the slopes calculated from the first 10 min of a titration of the 6 nt U substrate (0 - 1000 nM) performed in the presence of 75 nM nsp15 (see Supplementary Figure S2A and Experimental Procedures). **B.** Scheme describing the reaction steps during the screen. **C.** Residual activity of nsp15 in each sample well containing a compound for the low (1.25 μM) and high (3.75 μM) concentration of compounds. **D.** Comparison of residual activities of sample wells between low and high concentration. **E.** Comparison of Z-scores of each compound between low and high concentration.

We performed the screen with 75 nM nsp15 enzyme and 500 nM substrate, which produced reactions that were linear over the 15 min of reaction monitoring, against a library of over 5000 commercial compounds. These compounds were dispensed in DMSO over 24 384-well plates, and we tested two concentrations: 1.25 μM (low) and 3.75 μM (high). For the screen, we dispensed the nsp15 enzyme and incubated it with the drugs at room temperature (RT) for 10 min, after which we dispensed the substrate and started monitoring Cy5 fluorescence every minute for 15 minutes (Figure 3B and Supplementary Figure S2B). We included extra wells in every plate to control for any change in enzyme kinetics that could affect the identification of inhibitors (Supplementary Figure S2B). The Z’ factor was calculated for each plate, and the average Z’ factor for the whole screen was 0.8 indicating a high-quality screen (Supplementary Figure S2C). One plate of the high compound concentration had a Z’ factor <0.5 and was omitted from analysis.

Reaction slopes in the presence of each compound were calculated and normalised to the DMSO-containing positive controls (control reactions) on each plate (Figure 3C and Supplementary Figure S2B). We did not find any inhibitors that showed residual activities <0.7 in both concentrations, which was surprising given the results of similar screens from our accompanying manuscripts. Therefore, we considered compounds that had residual activities <0.9 and had a Z-score <-4 (Figure 3E and Supplementary Figure S2D). We also considered hits that decreased activity further in the high concentration than the low concentration, even if both residual activities were not below our cut-offs.

### Validation of Hits

We selected 17 compounds of the top hits from the screen for further analysis (Supplementary Table S3). We performed a test to determine if any of the 17 compounds quenched Cy5 fluorescence. Four compounds (in red) quenched Cy5 fluorescence in a concentration-dependent manner (Supplementary Figure S3). We then assessed inhibition of nsp15 endoribonuclease activity with the 12 remaining compounds using the plate reader (Supplementary Figure S4). Only NSC95397 and BMS-1166 compounds were able to inhibit nsp15 endoribonuclease activity at 10 μM (Figure 4A and Supplementary Figure S4). To explore inhibition by the top hit compounds further, we returned to the gel-based assay and assessed nsp15 endoribonuclease activity using the 16 nt substrate (Figure 4B). Again, NSC95397 (drug 10) inhibited nsp15 endoribonuclease. BMS-1166 did not inhibit the enzyme even at 100 μM in the gel-based assays. GNF-PF-3777 (drug 6) had been chosen for its high score as a putative activator of nsp15 (Supplementary Table S3) but had no effect even at 100 μM in the gel-based assays (Figure 4B). WEHI-539 hydrochloride (drug 1), selected from the screen as a putative nsp15 inhibitor and later found to quench Cy5 fluorescence (Supplementary Figure S3), enhanced nsp15 endoribonuclease activity in the gel-based assay (Figure 4B).

**Figure 4.**
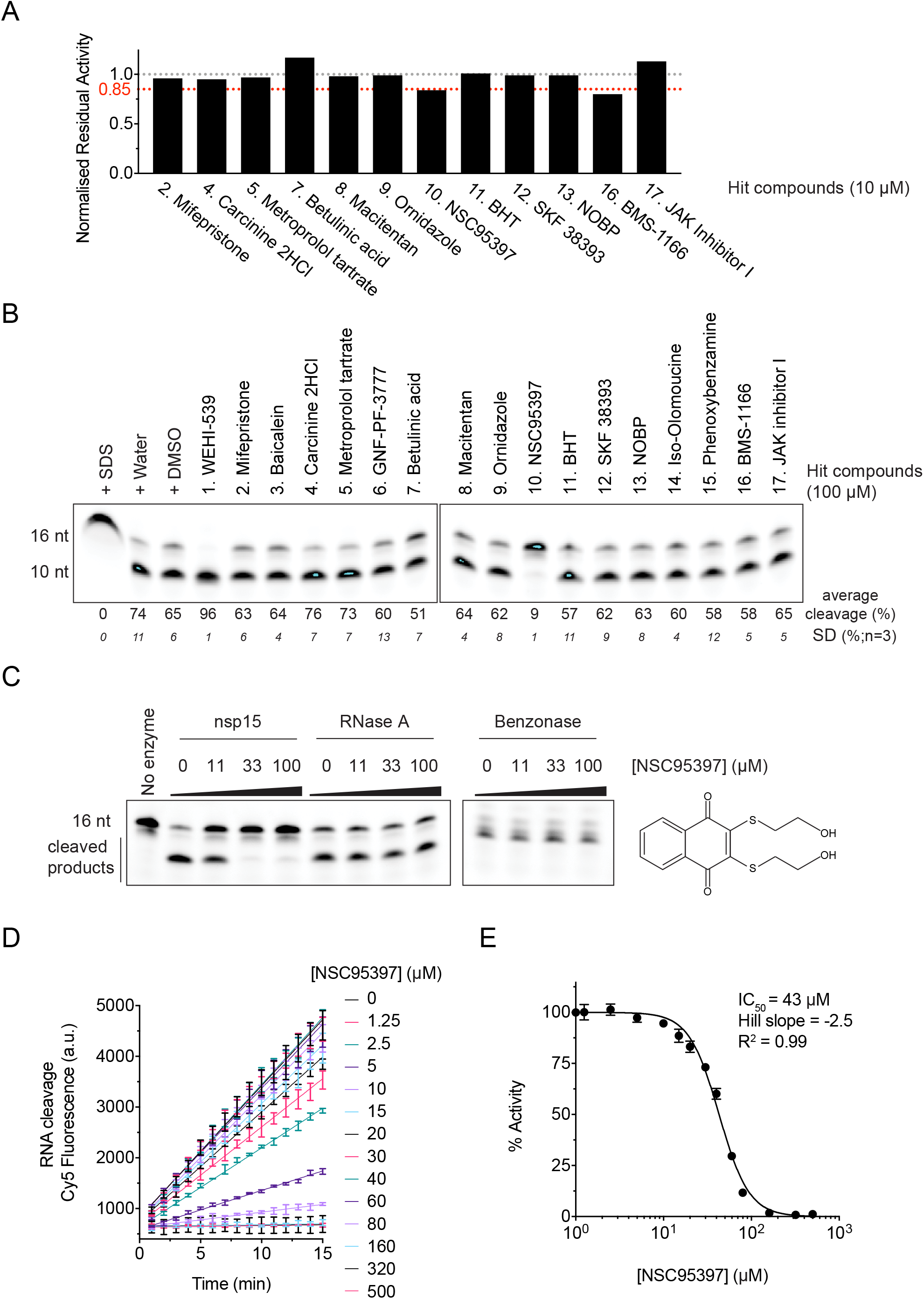
NSC95397 inhibits SARS-CoV-2 nsp15 endoribonuclease activity in vitro. **A.** Normalised residual activity of nuclease reactions monitoring cleavage of the 6 nt U substrate (500 nM) in solution using a Spark Multimode microplate reader (Tecan) in the presence of 75 nM of nsp15 and 10 μM of each of the 12 non-quenching selected screen hits (Supplementary Figure S3). Residual activities were calculated for each compound from experiment shown in Supplementary Figure S4, and normalised to control reaction. **B**. Nuclease reactions containing 500 nM nsp15 enzyme and 1 μM 16 nt U substrate in the presence of 100 μM of each of the 17 selected screen hits. Reactions were performed for 20 min at 30°C and resolved in a denaturing TBE-urea denaturing polyacrylamide gel. Control lanes include enzyme denaturation by SDS and addition of water or DMSO, to mimic addition of drugs diluted either in water or DMSO. **C**. Nuclease reactions containing 500 nM nsp15 fusion, 1 pg/μl RNase A or 25 mU/μl Pierce Universal Nuclease (benzonase) and 1 μM 16 nt U substrate in the presence of 0, 11, 33 or 100 μM NSC95397 inhibitor. Reactions were performed as in B. **D.** Titration of NSC95397 inhibitor (0 – 100 μM) in the presence of 75 nM nsp15 enzyme and 500 nM 6 nt U substrate and fluorescence quantified in a Spark Multimode microplate reader (Tecan) at RT every min for 15 min. **E.** Dose-response curves and IC50 values of NSC95397 for SARS-CoV-2 nsp15. IC50 values were calculated as described in Experimental Procedures.

To study the specificity of NSC95397 towards nsp15, we tested whether RNase A and benzonase activities were inhibited by NSC95397. Again, NSC95397 inhibited nsp15 endoribonuclease activity, but did not inhibit RNase A or benzonase, indicating that NSC95397 is not a general nuclease inhibitor (Figure 4C).

To determine the range of concentration of NSC95397 able to inhibit nsp15 endoribonuclease, we titrated NSC95397 and estimated the IC_50_ (43 μM) of NSC95397 inhibition of nsp15 endoribonuclease (Figure 4D and 4E).

### Effects of NSC95397 on SARS-CoV-2 growth in VERO E6 cells

Previous work has suggested that nsp15 activity plays a role in host recognition and response and is dispensable for viral replication and growth [21, 25, 27]. However, another study found that nsp15 inhibitors reduced SARS-CoV-1 viral growth in VERO E6 cells [31]. The compounds they used inhibited nsp15 endoribonuclease activity *in vitro*, but they also inhibited other ribonucleases. So, these results could be explained by inhibition of nsp15 or other viral or host RNases. Since the compound from our screen, NSC95397, selectively inhibited nsp15 and not RNase A or benzonase, we used NSC95397 to determine whether specific inhibition of nsp15 reduces SARS-CoV-2 viral growth in VERO E6 cells.

We infected VERO E6 cells with SARS-CoV-2 in the presence of a range of concentrations of NSC95397 and quantified viral infected area (Figure 5A and 5B). NSC95397 killed VERO E6 cells at concentrations above 10 μM, consistent with previous reports that NSC95397 is an inhibitor of the Cdc25 protein phosphatase and multiple protein kinases [32–35]. This made it more difficult to assess viral growth. With the aim of finding a concentration of NSC95397 that inhibited viral growth without killing the cells, we performed a second titration at lower NSC95397 concentrations (Figure 5C and 5D). Cell survival was much less impacted, but viral growth was not inhibited by NSC95397 (Figure 5C and 5D).

**Figure 5.**
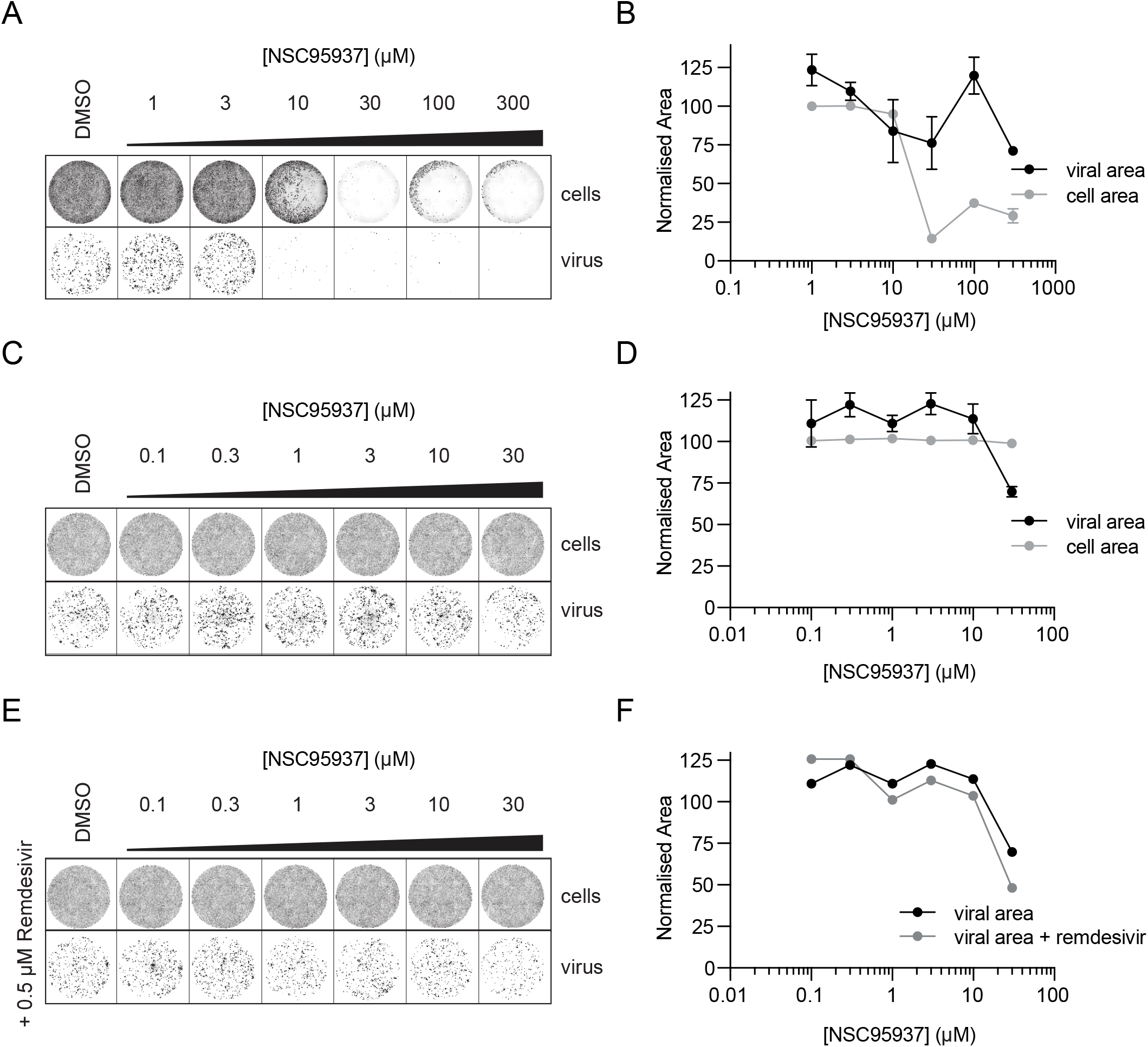
NSC95397 does not inhibit SARS-CoV-2 growth in VERO cells. **A.** SARS-CoV-2 infectivity assays in the presence of NSC95397. DNA in cells was stained with DRAQ5, and virus was immunostained with anti-N antibody. **B.** Quantification of normalised viral area (black) and cell area (gray) +/-SD. Both viral and cell area was normalised to control condition (DMSO) and, viral area was also normalised to cell area (see Experimental Procedures for details). **C.** SARS-CoV-2 infectivity assays as in A with lower concentrations of NSC95397. **D**. Quantification of C as in B. **E**. SARS-CoV-2 infectivity assays as in C in the presence of 0.5 μM remdesivir. **F**. Quantification of E as in B, note that black control line is quantification of control experiment in C.

Mutation of the exoribonuclease activity of nsp14 has been shown to sensitise SARS-CoV-1 to the antiviral agent remdesivir [36]. To explore if inhibition of the nsp15 endoribonuclease activity also synergised with remdesivir antiviral activity, we performed infectivity assays in the presence of both NSC95397 and remdesivir. However, we did not observe any additive effect of these two drugs on SARS-CoV-2 infectivity in VERO E6 cells (Figure 5E and 5F).

## Discussion

We have purified SARS-CoV-2 nsp15 enzyme and optimised fluorescent biochemical endoribonuclease (EndoU) assays to screen a custom chemical library containing over 5000 characterised commercial compounds. We identified NSC95397 as a novel nsp15 EndoU inhibitor *in vitro*. The role of nsp15 endoribonuclease during SARS-CoV-2 proliferation is still unclear [18, 20, 21, 25, 37]. Coronaviruses with mutated nsp15 are viable in viral replication assays yet have severe growth defects in immune specialised cell models such as macrophages and in immune proficient mouse models, indicating an essential role for nsp15 in mediating the evasion of the host immune system [21, 26, 27, 38, 39]. Indeed, nsp15 has been shown to antagonise the production of interferon beta, and thus its inhibition might also lead to increased interferon production [40]. Macrophages and other immune cells can become infected with SARS-CoV-2 in human patients, and lung macrophage dysfunction has been associated with severe COVID-19, consistent with a key role for macrophages in the early immune response to combat SARS-CoV-2 infections [41–43]. Therefore, the use of nsp15 inhibitors would appear to be an attractive strategy to boost immune recognition of the virus at early stages of infection.

Although NSC95397 inhibited nsp15 activity *in vitro*, it did not impair the proliferation of SARS-CoV-2 in VERO E6 cells, which are a commonly used host cell for viral growth under laboratory conditions. However, VERO E6 are derived from the Green Monkey *Chlorocebus sabaeus*, and it will be important in future studies to establish better cellular models with which to assess the importance of nsp15 for the evasion of host immune responses during SARS-CoV-2 infection of appropriate human cells.

Being an endoribonuclease, unrestricted nsp15 activity might lead to unwanted cleavage of the viral genome, so regulation of its activity is likely to be important for balancing its immune evasion function with viral genome integrity. One level of regulation of nsp15 activity is at the substrate level where nsp15 shows differential activity towards sequences surrounding the targeted uridylate, as well as RNA secondary structures and modifications such as 2’-O-methyl groups [18, 44]. The formation of the nsp15 active hexamer could be another level of regulation [45]. Because nsp15 is only active as a hexamer, it is likely that nsp15 activity is restricted to later steps of infection when enough nsp15 has been translated to support hexamer formation. Our results show that nsp15 activity is stimulated at high substrate concentrations, indicating that nsp15 is an allosteric enzyme, which confirms predictions based on structural cryo-EM studies of the SARS-CoV-2 nsp15 hexamer [23] (Figure 2, 3A and Supplementary Figure S2A). This allosteric behaviour might serve as an additional control to limit nsp15 EndoU activity and locally protect the viral RNA. Thus, nsp15 activation might occur at later times post infection, where the virus has produced enough nsp15 protein and substrate.

Accordingly, in our screen we used high enzyme and substrate concentrations (75 nM nsp15 and 500 nM substrate). However, this high enzyme concentration combined with high substrate competition (for potential competitive inhibitors) might explain why we did not detect many compounds able to inhibit nsp15 EndoU activity and our hit compounds only reduced nsp15 activity by 10-20% in the original conditions of the screen. We and our colleagues were able to detect hits for the other SARS-CoV-2 enzymes using the same drug library and general screening strategy (see accompanying manuscripts) and, therefore, it would be advisable for future studies to use higher compound to enzyme or substrate ratios to improve detection of nsp15 inhibitors.

We found that NSC95397 is an inhibitor of nsp15 but not a general nuclease inhibitor *in vitro*. We looked for compounds previously shown or predicted to inhibit nsp15 in our library and found that, although none passed our cut-off, some of them also inhibited nsp15 to some degree in our screen (Supplementary Table S4) [31, 46–48]. Previous studies have found nsp15 to be sensitive to RNase A inhibitors [31], so, interestingly, our data would suggest that NSC95397 inhibits nsp15 through a different mechanism (Figure 4C). In addition, though our infectivity assays using VERO E6 cells did not find NSC95397 to be effective in inhibiting SARS-CoV-2 viral growth, RNase A inhibitors did reduce SARS-CoV-1 viral growth in VERO cells [31]. This could be due to RNase A inhibitors also inhibiting the other nuclease in coronaviruses, nsp14, or it could be that the higher viral load and later timepoint of infection used in that study compared to our infectivity assays could have been more sensitive to nsp15 inhibition. In agreement with a late infection role for nsp15, nsp15 can specifically cleave the polyuridine tracks of the later transcribed negative-strand RNA strand of the virus, which could lead to reducing the activation of cytosolic immune sensors [38]. Future studies could assess the effects of NSC95397 with late infection conditions, in appropriate human immune cells or in mouse models, which, as discussed above, are sensitive to nsp15 mutations. In addition, our assays showed high host cell toxicity in response to NSC95397, so it would also be interesting to test fluorinated and hydroxylated derivatives of NSC95397, which have been proposed to be less toxic [49].

Because of its role in immune evasion, treatment with nsp15 inhibitors could be useful early in SARS-CoV-2 infection to help elicit a robust immune response to clear the virus. As an alternative strategy, nsp15 inhibitors could be tested in combination with other treatments to screen for synthetic anti-viral effects. Our colleagues have identified other SARS-CoV-2 enzyme inhibitors (see accompanying manuscripts), and it would be interesting to test multiple compounds in a “cocktail” approach to treating COVID-19.

## Experimental Procedures

### Expression constructs

We constructed 4 plasmids to express different tagging combinations of nsp15 in *Escherichia coli* (*E.coli*) (Supplementary Table S1 and S2). SARS-CoV-2 nsp15 was codon optimised for bacterial expression (Genbank MN908947.3) and cloned into BamHI-NotI MCS of pMEXCb vector (https://mrcppu-covid.bio/cdna-clones/134146) to create plasmid SARS-CoV-2 MBP-nsp15 (DU67734). Plasmid SARS-CoV-2 14His-SUMO-nsp15 (DU70489) was built by PCR with oligos 9142/9143 from DU67734 and Gibson assembly into K27SUMO vector [50]. Plasmid SARS-CoV-2 6His-nsp15 (DU70490) was built by PCR with oligos 9156/9157 from DU67734 followed by PCR digestion with NcoI and XhoI and ligation into pET28C. Plasmid SARS-CoV-2 His-nsp15 (H234A H249A) (DU70491) was built by PCR with oligos 9150/9151 from DU70490.

### Expression and purification of SARS-CoV-2 nsp15 in E. coli

Rosetta™ (DE3) pLysS cells (Novagen) (F- ompT hsdSB (rB- mB-) gal dcm (DE3) pLysSRARE (CamR) were transformed with plasmid DU70490 to express 6His-nsp15. Transformant colonies were inoculated into a 400 ml LB / chloramphenicol (35 μg/ml) / kanamycin (50 μg/ml) and grown overnight at 37°C with shaking at 200 rpm. The next morning, the culture was mixed with 3600 ml of LB / chloramphenicol (35 μg/ml) / kanamycin (50 μg/ml) and further grown at 37°C until OD_600_ reached 0.8. Protein expression was induced by addition of 0.4 mM IPTG, and the culture was shaken overnight at 16°C. Cells were harvested by centrifugation at 5000 rpm for 10 min in a JLA-9.1000 rotor (Beckman). The bacterial pellet was resuspended in 20 ml lysis buffer (50 mM Tris-HCl (pH 8), 0.5 M NaCl, 10 mM imidazole, 0.5 mM TCEP, Roche protease inhibitor tablets) with 500 μg / ml lysozyme, then incubated at 4°C for 0.5 h with rotation. Subsequently, the sample was sonicated twice for 90 s (15 s on, 30 s off) at 40% on a Branson Digital Sonifier. After centrifugation at 15,000 rpm at 4°C for 0.5 h in an JA-30.50 rotor (Beckman), the obtained soluble extract was mixed with 2 ml slurry Ni-NTA beads (30210, QIAGEN), incubated at 4°C for 2 h with rotation. Beads were recovered in a disposable gravity flow column and washed with 150 ml of lysis buffer then 20 ml lysis buffer containing 10 mM MgCl_2_ and 2 mM ATP (to remove bacterial chaperones) then 20 ml lysis buffer. Proteins were eluted with 5 ml lysis buffer containing 0.4 M imidazole. The eluate was loaded onto a 120 ml Superdex 200 column in Gel filtration buffer (50 mM Tris-HCl (pH 8), 0.15 M NaCl, 0.5 mM TCEP). Nsp15-containing fractions were pooled, concentrated to 0.6 mg/ml by ultrafiltration using Amicon Ultra centrifugal unit (30 k MWCO; MERCK), aliquoted and snap frozen.

The 6His-tagged H234A H249A nsp15 mutant was purified as described above. 14His-SUMO-nsp15 was purified as for 6His-nsp15, except that we attempted to remove the 14His-SUMO tag by overnight Ulp1 cleavage at 4°C before the gel filtration step. Proteolytic cleavage was partial, and the resultant untagged nsp15 was subsequently separated from 14His-SUMO-nsp15 on a 120 ml Superdex 200 column equilibrated in gel filtration buffer (50 mM Tris-HCl (pH 8), 0.15 M NaCl, 0.5 mM TCEP).

For purification of MBP-nsp15, Rosetta™ (DE3) pLysS cells were transformed with plasmid DU67734 (pMEX3Cb SARS-CoV2 (2019-nCoV) NSP15). Transformant colonies were inoculated into 200 ml LB / chloramphenicol (35 μg/ml) / ampicillin (100 μg/ml) and grown at 37°C with shaking at 200 rpm. After ~10 h growth, 100 ml of the culture was mixed with 900 ml of LB / chloramphenicol (35 μg/ml) / ampicillin (100 μg/ml) and grown at 37°C until OD_600_ reached 0.8. The protein expression was induced by addition of 0.2 mM IPTG, and the culture was shaken overnight at 18°C. Cells were harvested by centrifugation at 5000 rpm for 10 min in a JLA-9.1000 rotor (Beckman). The bacterial pellet was resuspended in 20 ml lysis buffer (50 mM Tris-HCl (pH 8), 0.5 M NaCl, 0.5 mM TCEP, Roche protease inhibitor tablets) and lysed as described for 6His-nsp15. The clarified extract was mixed with 2 ml amylose beads and incubated at 4°C for 2 h with rotation. Beads were recovered and washed as for 6His-nsp15. Proteins were eluted with 5 ml lysis buffer containing 10 mM maltose. The eluate was loaded onto a 120 ml Superdex 200 column in Gel filtration buffer (50 mM Tris-HCl (pH 8), 0.15 M NaCl, 0.5 mM TCEP). Nsp15-containing fractions were pooled, concentrated by ultrafiltration using Amicon Ultra centrifugal unit (30 k MWCO; MERCK), aliquoted and snap frozen.

### Expression and purification of SARS-CoV-2 nsp15 in insect cells

The plasmid SARS-CoV-2 3xFlag-His-nsp5CS-nsp15 (Addgene ID: 169166) was used to express 3xFlag-His-nsp5CS-nsp15 in baculovirus-infected insect cells (Supplementary Table S1 and S2). The coding sequence of SARS-CoV-2 nsp15 (NCBI reference sequence NC_045512.2) was codon-optimised for *S. frugiperda* and synthesised (GeneArt, Thermo Fisher Scientific) (codon optimised sequence can be found in supplementary information). Nsp15 was subcloned into a modified biGBac pBIG1a vector containing a pLIB-derived polyhedrin expression cassette [51] to contain an N-terminal 3xFlag-His6 tag (sequence: MDYKDHDGDYKDHDIDYKDDDDKGSHHHHHHSAVLQ-nsp15). Baculoviruses were generated and amplified in Sf9 cells (Thermo Fisher Scientific) using the EMBacY baculoviral genome (Trowitzsch et al., J Struct Biol, 2010). For protein expression Sf9 cells were infected with baculovirus, collected 48 hours after infection, flash-frozen, and stored at −70°C.Cell pellets were resuspended in pulldown buffer (30 mM HEPES pH 7.6, 250 mM sodium chloride, 5 mM magnesium acetate, 10% glycerol, 0.02% NP-40 substitute, 1 mM DTT) supplemented with protease inhibitors (Roche Complete Ultra tablets, 1 mM AEBSF, 10 μg/ml pepstatin A, 10 μg/ml leupeptin) and lysed with a dounce homogenizer. The protein was purified from the cleared lysate by affinity to Anti-FLAG M2 Affinity gel (Sigma-Aldrich) and eluted with pulldown buffer containing 0.1 mg/ml 3xFlag peptide. Eluate was further purified by gel filtration as described for the bacterially expressed proteins.

### SARS-CoV-2 nsp15 endoribonuclease assays

A 16 nt 5’ Cy3-single stranded RNA (ssRNA) substrate (16 nt substrate) was used to monitor the nsp15 uridine-dependent endoribonuclease activity in gel-based assays (Supplementary Table S2). A 6 nt 5’ Cy5 and 3’ BHQ650 quencher ssRNA substrate (6 nt substrate) was used to quantify nsp15 uridine-dependent endoribonuclease activity in gel-based assays (Figure 2B and 2C) and in solution using a Spark Multimode microplate reader (Tecan).

The assay, with either substrate, was performed by incubating the enzyme and the substrate at room temperature in total 20 μl in nsp15 reaction buffer (50 mM Tris-HCl pH 7.5, 50 mM NaCl, 10 mM MnCl_2_, 5 mM MgCl_2_, 0.1 mg/ml BSA, 0.02 % Tween-20, 10 % glycerol and 0.5 mM TCEP). Specific enzyme and substrate concentrations as well as duration of the reaction is indicated in the figure legends for each experiment.

### High-throughput kinetic endoribonuclease screen

High-throughput screen was performed using a custom collection of over 5000 compounds from commercial sources (Sigma, Selleck, Enzo, Tocris, Calbiochem, and Symansis). 2.5 or 7.5 nl of a 10 mM stock of the compounds dissolved in DMSO were arrayed and dispensed into square flat-bottom black 384-well plates containing 1 μl DMSO / well using an Echo 550 (Labcyte), before being sealed and stored at −80°C.

The day of the screen, plates were initially moved from −80°C to 4°C, then moved to room temperature for at least 30 min prior to the screen. Plates were centrifuged and desealed just prior to dispensing 10 μl of 2X enzyme mix (150 nM nsp15, 50 mM Tris-HCl pH 7.5, 50 mM NaCl, 10 mM MnCl2, 5 mM MgCl_2_, 0.1 mg/ml BSA, 0.02% Tween20, 10% glycerol, 0.5 mM TCEP) using a XRD-384 Reagent Dispenser (FluidX Ltd.) or hand-pippetting control columns (Figure 3B and Supplementary Figure S2B). After 10 min, 10 μl of 2x substrate mix (1000 nM 6 nt U substrate in same buffer as enzyme mix) was dispensed and plates were centrifuged. 2 min after dispensing substrate mix, plates were read with a Spark Multimode microplate reader (Tecan) with the following settings: Excitation 645 nm (+/-10), Emission 675 nm (+/-10), Gain 125, 10 flashes, Z position of 17500, every min for 15 min.

### Screen data analysis

The slope of each reaction was determined by linear regression. Residual activity was then calculated by dividing residual activity in the presence of each compound by the median of the control wells without drugs of each plate. Z-scores were calculated using the standard deviation and median of residual activities for each concentration. Plate 24 of the high concentration was omitted from analysis. Z’ factors were calculated for each plate to determine screen quality according to Zhang et al. [52].

### K_M_ and IC_50_ calculation

First, we calculated the slope of the reaction in the presence of 75 nM nsp15 and a titration of the 6 nt U substrate (0 – 1000 nM) considering the first 10 min of the reactions by linear regression. Slopes were then used to calculate K_half_ and V_MAX_ by non-linear fitting to an Allosteric sigmoidal equation using GraphPad Prism.

Similarly, we calculated the slope of the reaction in the presence of 75 nM nsp15, 6 nt U substrate (500 nM) and a titration of NSC95397 considering the first 15 min of the reactions by linear regression. Slopes were then used to calculate percentage of activity relative to the slope obtained with 0 μM NSC95397. Percent activity for each log_10_ of the concentration of NSC95397 (μM) were then used to estimate the IC_50_ and Hill slopes of NSC95397 for nsp15 endoribonuclease using GraphPad Prism.

### SARS-CoV-2 infectivity assays

Infectivity assays were performed as in the accompanying manuscripts. Briefly, VERO E6 cells were grown in 96 well imaging plates and transfected with individual gapmers at 0.5 μM using Lipofectamine 2000 (Thermofisher). 6 h post transfection, the media was replaced and infected with SARS-CoV-2 at an MOI of 0.5 PFU/cell and, simultaneously, a titration of NSC95397 was added to different wells with or without the addition of 0.5 μM remdesivir. 22 h post-infection, cells were fixed, permeabilised and stained for SARS-CoV-2 N-protein using Alexa488-labelled-CR3009 antibody [53] and nuclei using DRAQ5 (647 nm wavelength). Imaging was performed using a 5x lens in an Opera Phenix high content screening microscope (Perkin Elmer) equipped with the Harmony software. The whole well area was delineated, and area of Alexa488/N protein and DRAQ5/DNA signals were determined for each well. The Alexa488/N intensities were normalised to DRAQ5/DNA and to vehicle treated samples.

## Data Availability Statement

All data associated with this paper will be deposited in FigShare (https://figshare.com/).

## Author Contributions

**Berta Canal:** Conceptualisation, Methodology, Validation, Formal analysis, Investigation, Resources, Writing – Original Draft, Writing – Review and Editing, Visualisation. **Ryo Fujisawa:** Conceptualisation, Methodology, Validation, Formal analysis, Investigation, Resources, Writing – Original Draft, Writing – Review and Editing, Visualisation. **Allison W. McClure:** Conceptualisation, Methodology, Validation, Formal analysis, Investigation, Resources, Writing – Original Draft, Writing – Review and Editing, Visualisation. **Tom Deegan:** Conceptualisation, Methodology, Validation, Formal analysis, Investigation, Resources, Writing – Review and Editing, Visualisation. **Mary Wu:** Methodology, Investigation, Resources. **Rachel Ulferts:** Methodology, Investigation. **Florian Weissmann:** Resources. **Lucy S. Drury:** Investigation, Resources. **Agustina P. Bertolin:** Resources. **Jingkun Zeng:** Resources, Software. **Rupert Beale:** Supervision. **Michael Howell:** Supervision. **Karim Labib:** Conceptualisation, Methodology, Writing – Review and Editing, Supervision. **John F.X Diffley:** Conceptualisation, Methodology, Writing – Review and Editing, Supervision, Project administration, Funding acquisition.

## Acknowledgements

We thank the Crick high-throughput screen (HTS) science technology platform (STP) for providing the chemical library and for help with screen design and analysis and are grateful to MRC Reagents and Services (https://mrcppureagents.dundee.ac.uk/) for providing DNA constructs for nsp15. We thank David McClure for help with screen analysis. We thank Anne Early for ordering supplies and hit compounds. This work was supported by the Francis Crick Institute, which receives its core funding from Cancer Research UK (FC001066), the UK Medical Research Council (FC001066), and the Wellcome Trust (FC001066). This work was also funded by a Wellcome Trust Senior Investigator Award (106252/Z/14/Z) to J.F.X.D. BC and FW have received funding from the European Union’s Horizon 2020 research and innovation programme under the Marie Skłodowska-Curie grant agreement Nos 895786 and 844211. JZ has received funding from a Ph.D. fellowship awarded by Boehringer Ingelheim Fonds. This work was also funded by the Medical Research Council (core grant MC_UU_12016/13 to KL), Cancer Research UK (Programme Grant C578/A24558 to KL) and the Wellcome Trust (reference 204678/Z/16/Z for a Sir Henry Wellcome Postdoctoral Fellowship to TD).

## Supplementary Material

**Supplementary Figure S1.**
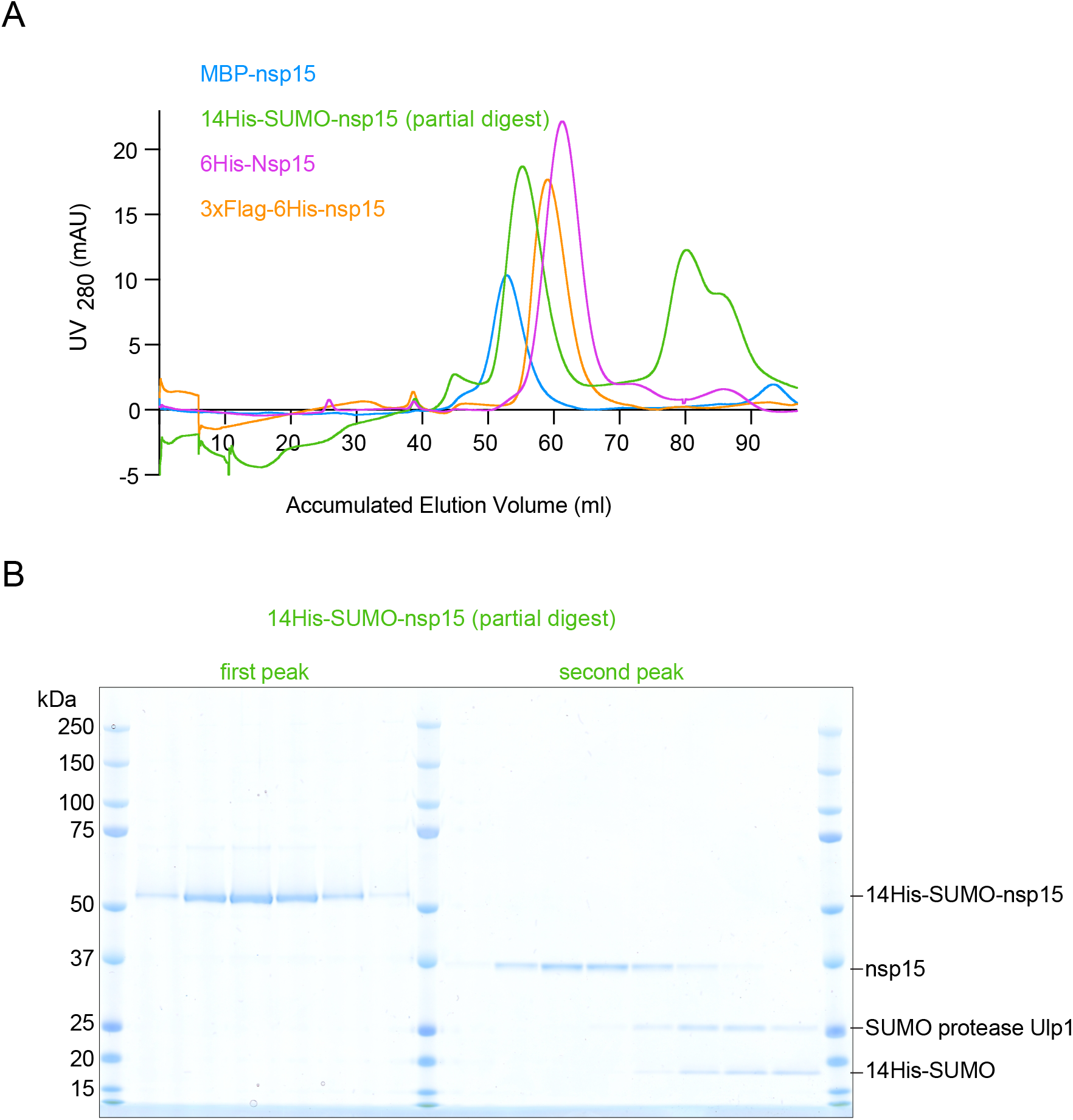
Size exclusion chromatography of purified SARS-CoV-2 nsp15. **A**. Each of the indicated protein preparations were loaded onto a 120 ml Superdex200 column, which has a void volume of ~40 ml. Approximate protein amounts as measured by UV_280_ are plotted versus accumulated elution volume. Note the presence of the later-eluting peak of the partially cleaved 14His-SUMO-nsp15 preparation, consistent with the untagged nsp15 existing as a smaller complex or monomer. **B**. Fractions from the size-exclusion chromatography of A for the partially digested 14His-SUMO-nsp15 were run on SDS-PAGE and stained with coomassie.

**Supplementary Figure S2.**
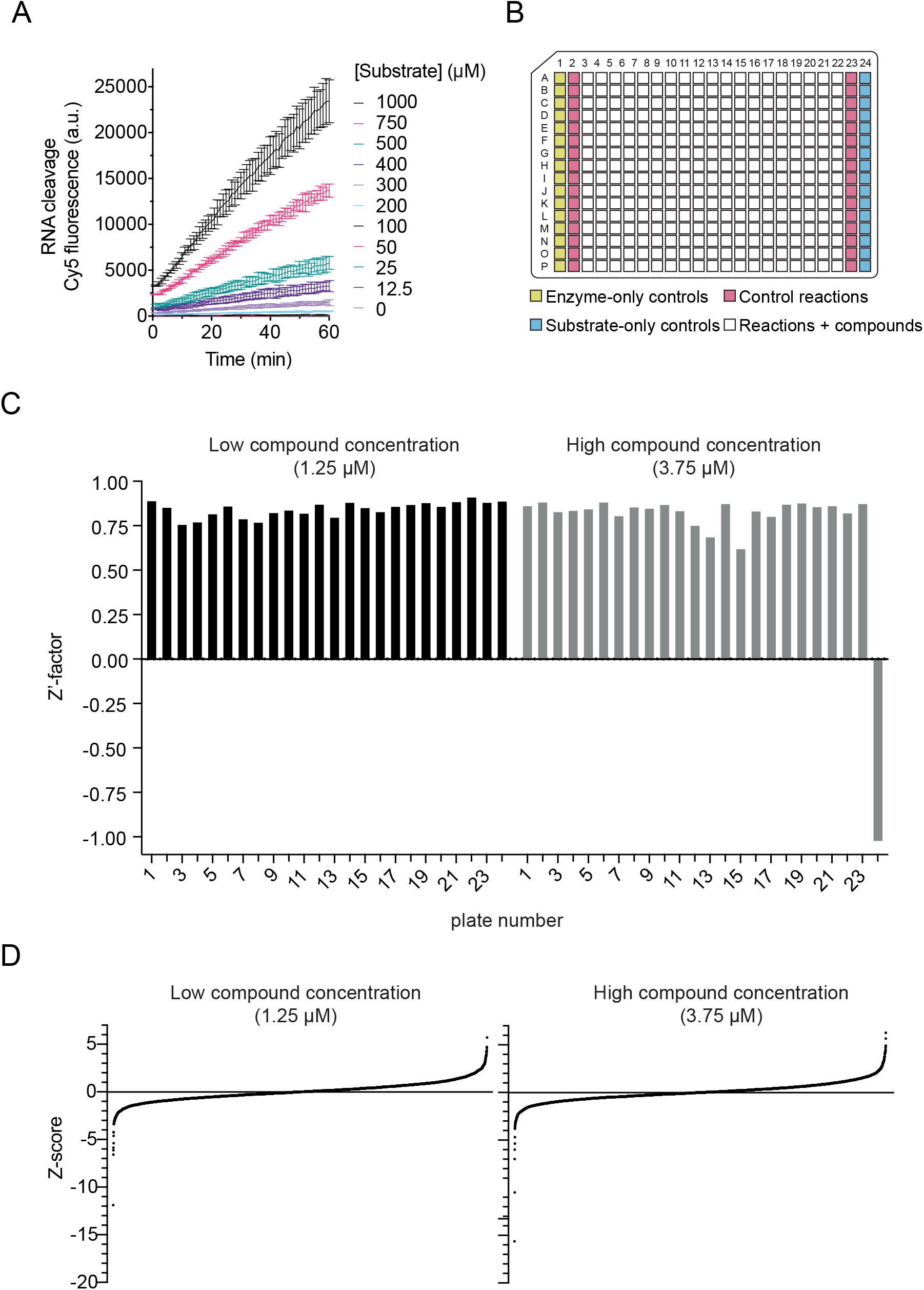
SARS-CoV-2 endoribonuclease nsp15 inhibitor screen design and results. **A.** Titration of the 6 nt U substrate (0 – 1000 nM) in the presence of 75 nM nsp15 enzyme. Fluorescence quantified in a Spark Multimode microplate reader (Tecan) at RT every min for 60 min. **B.** Schematic of the distribution of the controls (substrate-only, enzyme-only and control reactions) and of the compound reactions (reactions + compounds) in the plates of the screen. **C**. Z’ factor for each plate at both compound concentrations. **D**. Ordered Z-scores for sample wells at both concentrations.

**Supplementary Figure S3.**
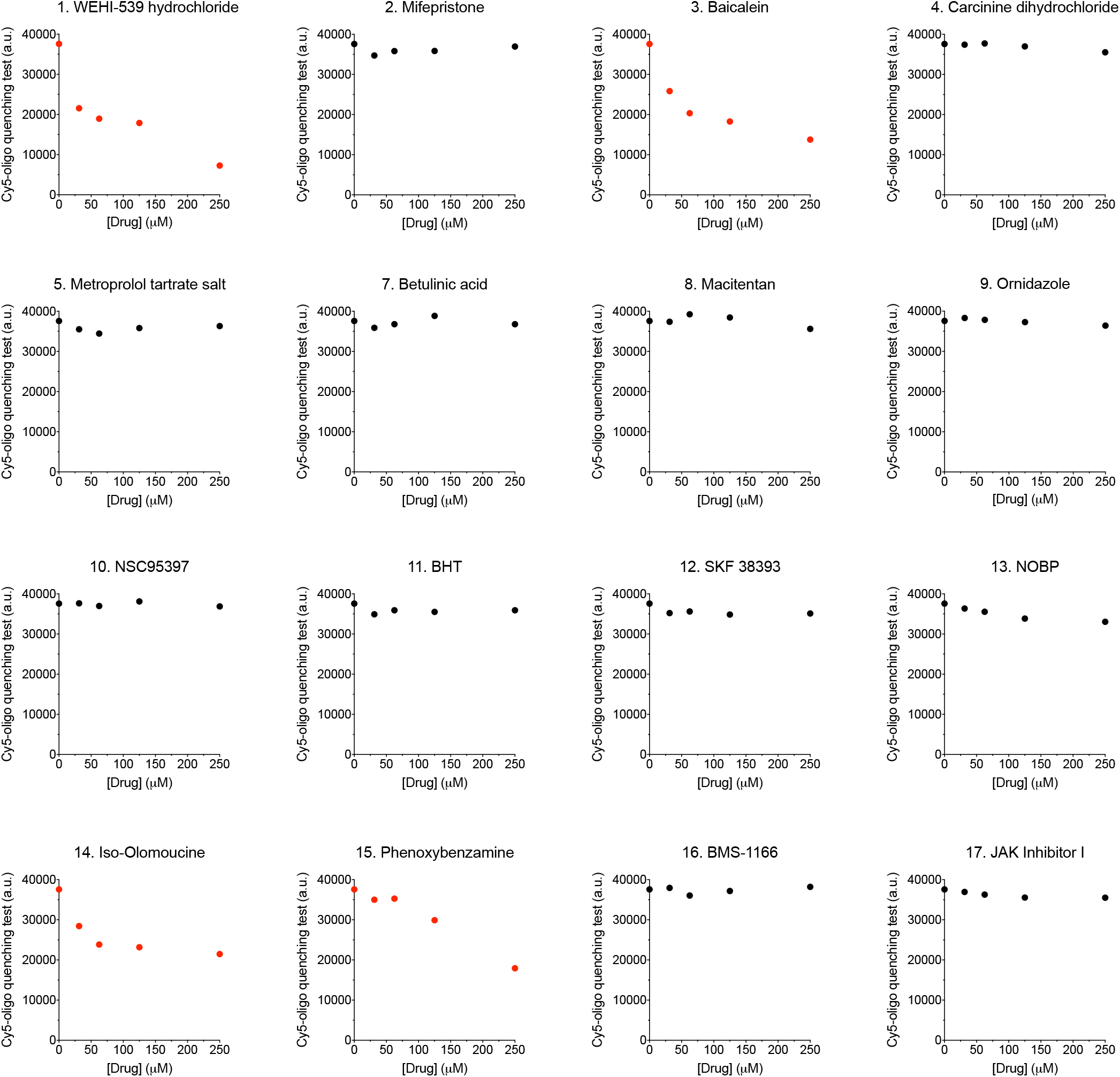
Fluorescence quenching test for screen hit compounds. Graphs show the fluorescence of a 35 nt Cy5-oligonucleotide (Supplementary Table S2) in the presence of a titration of the top hit compounds from the screen (except compound 6). In red, compounds that quench Cy5 fluorescence.

**Supplementary Figure S4.**
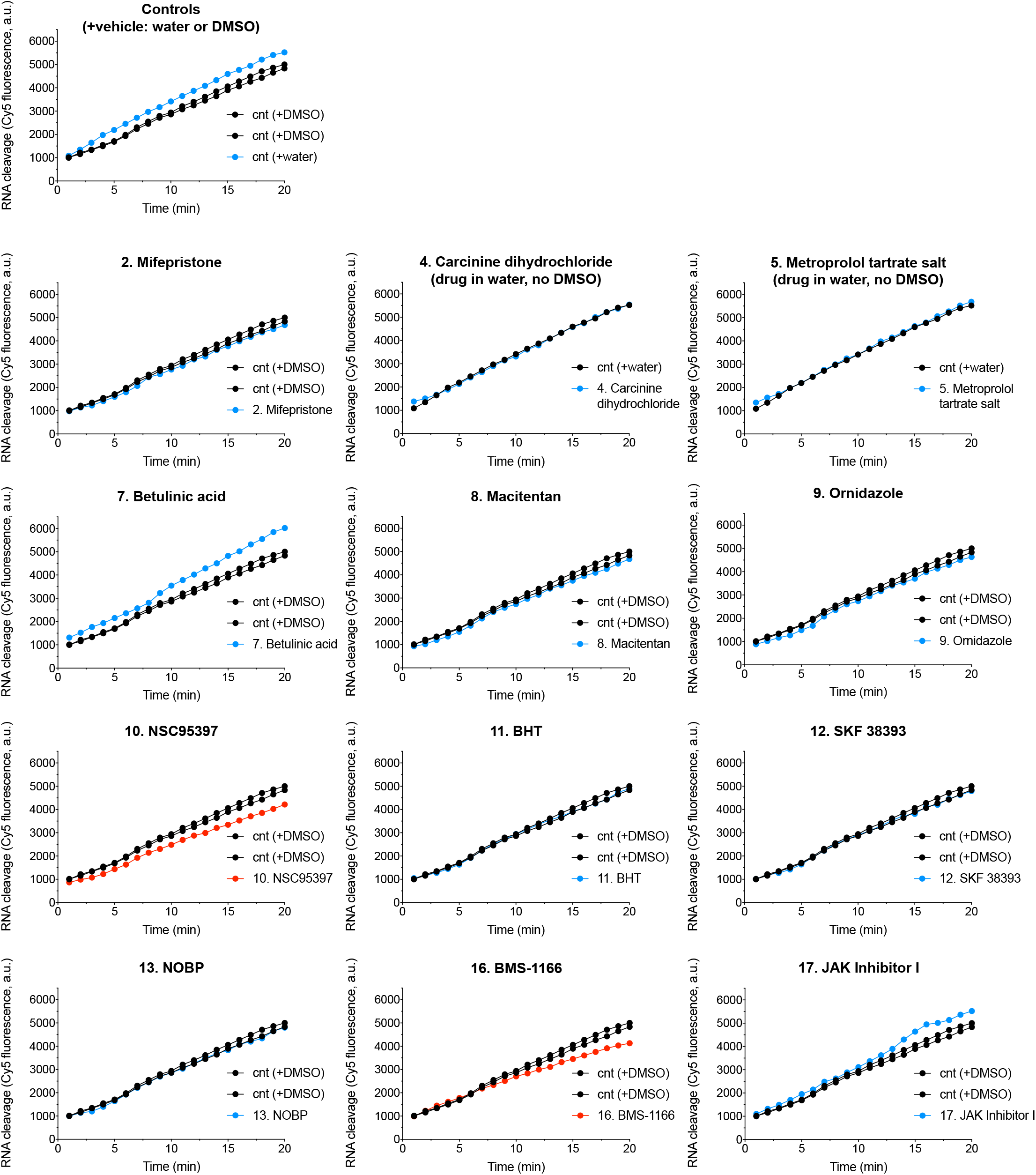
Validation of screen hit compounds in plate reader. Nuclease reactions in the presence of 75 nM nsp15, 500 nM 6 nt U substrate and 10 μM of the 12 non-quenching selected screen hits (Supplementary Figure S3, except 6). Up top graph shows three control reactions (2 +DMSO, 1 +water). The rest of the graphs show control reactions (in black) and the reactions in the presence of the different compounds (in blue -no inhibition-, and in red -inhibition-). Water control reactions are shown for compounds 4 and 5, that were dissolved in water. Residual activity for each compound was calculated from this experiment and shown in Figure 4A.

---

**Supplementary Table S1.** Cloning strategies.

| Supplementary Table S1. Cloning strategies |  |  |  |
| --- | --- | --- | --- |
| Expression | Construct ID | Available from | Cloning |
| Constructs for expression in *E. coli* |  |  |  |
| SARS-CoV-2 MBP-nsp15 | DU67734 | MRC PPU | Codon optimised SARS-CoV-2 nsp15 (Genbank MN908947.3) for bacterial expression, cloned into BamHI-NotI MCS of pMEXCb vector (https://mrcppu-covid.bio/cdna-clones/134146) |
| SARS-CoV-2 His-SUMO-nsp15 | DU70489 | MRC PPU | PCR with oligos 9142/9143 from DU67734, Gibson assembly into K27 vector |
| SARS-CoV-2 His-nsp15 | DU70490 | MRC PPU | PCR with oligos 9156/9157 from DU67734, Digest PCR fragment with NcoI and XhoI ligated into pET28C |
| SARS-CoV-2 His-nsp15 (H234A H249A) | DU70491 | MRC PPU | PCR with oligos 9150/9151 from DU70490 |
| Constructs for expression in baculovirus-infected Sf9 insect cells |  |  |  |
| SARS-CoV-2 3xFlag-His-nsp5CS-nsp15 | 169166 | Addgene | PCR with oligos FW701, FW702, FW758, FW759 from codon optimised nsp15 cloned into pBIG1a plasmid. |

**Supplementary Table S2.**
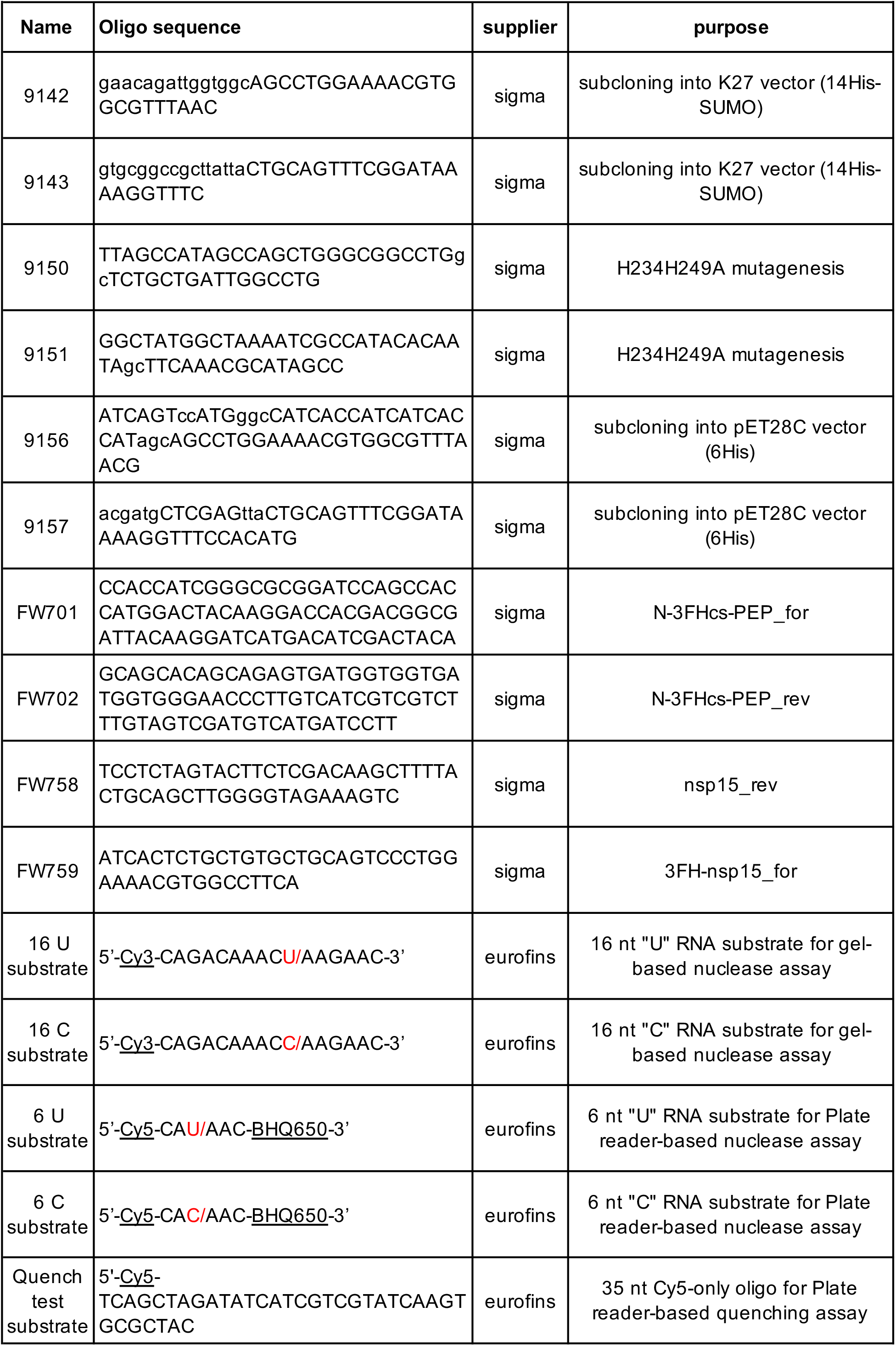
Cloning and substrate oligonucleotides.

**Supplementary Table S3.**
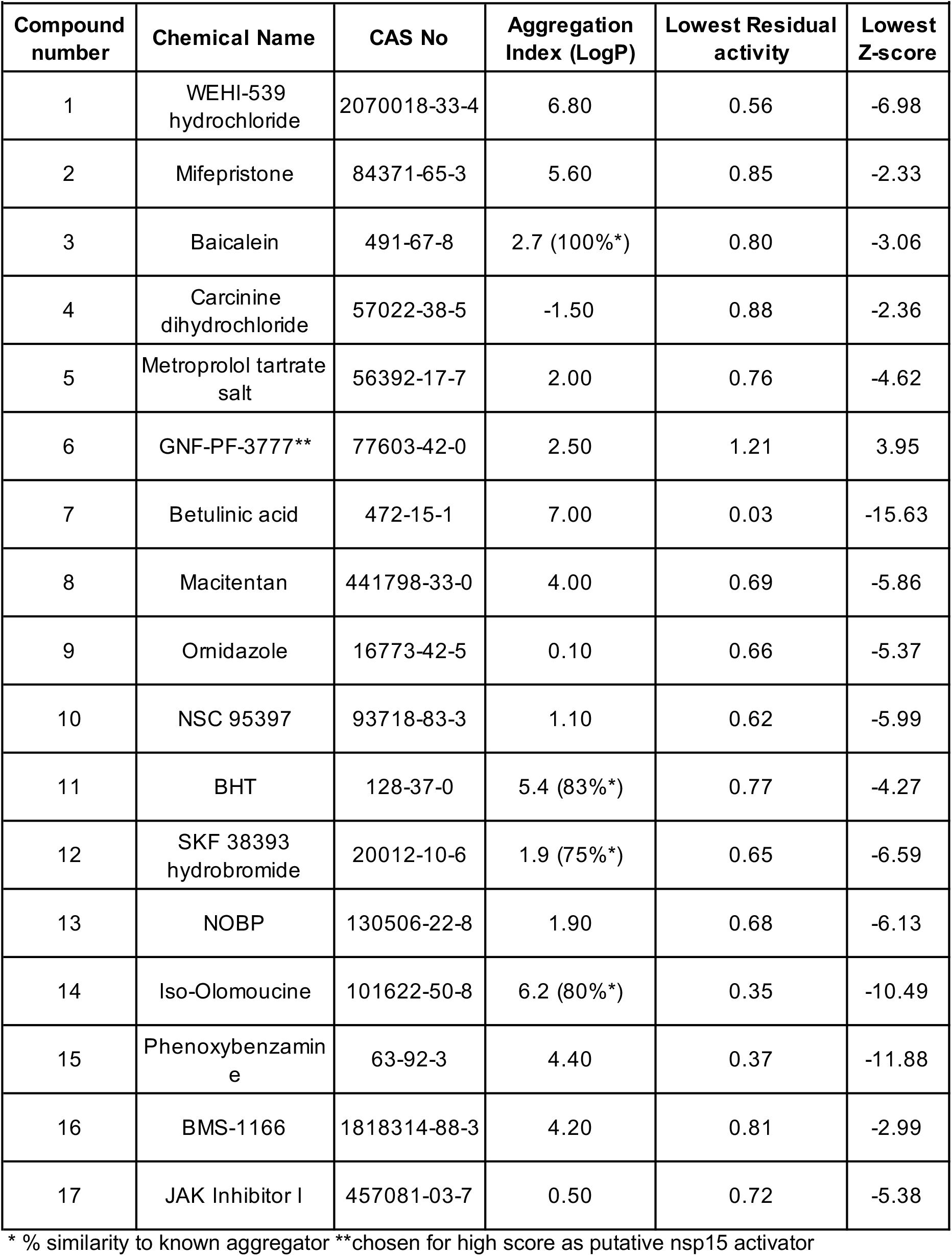
Top screen hit compounds.

**Supplementary Table S4.**
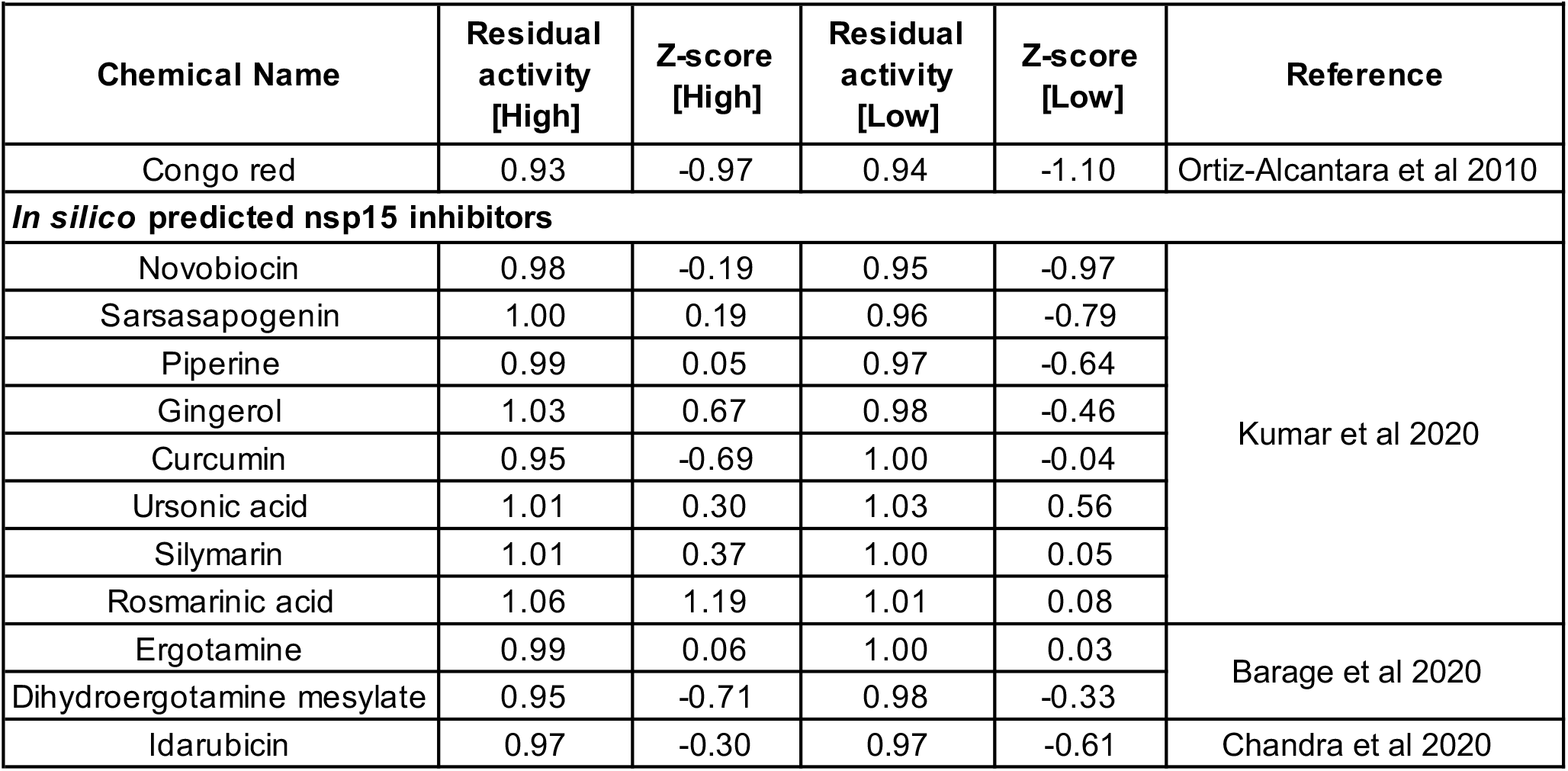
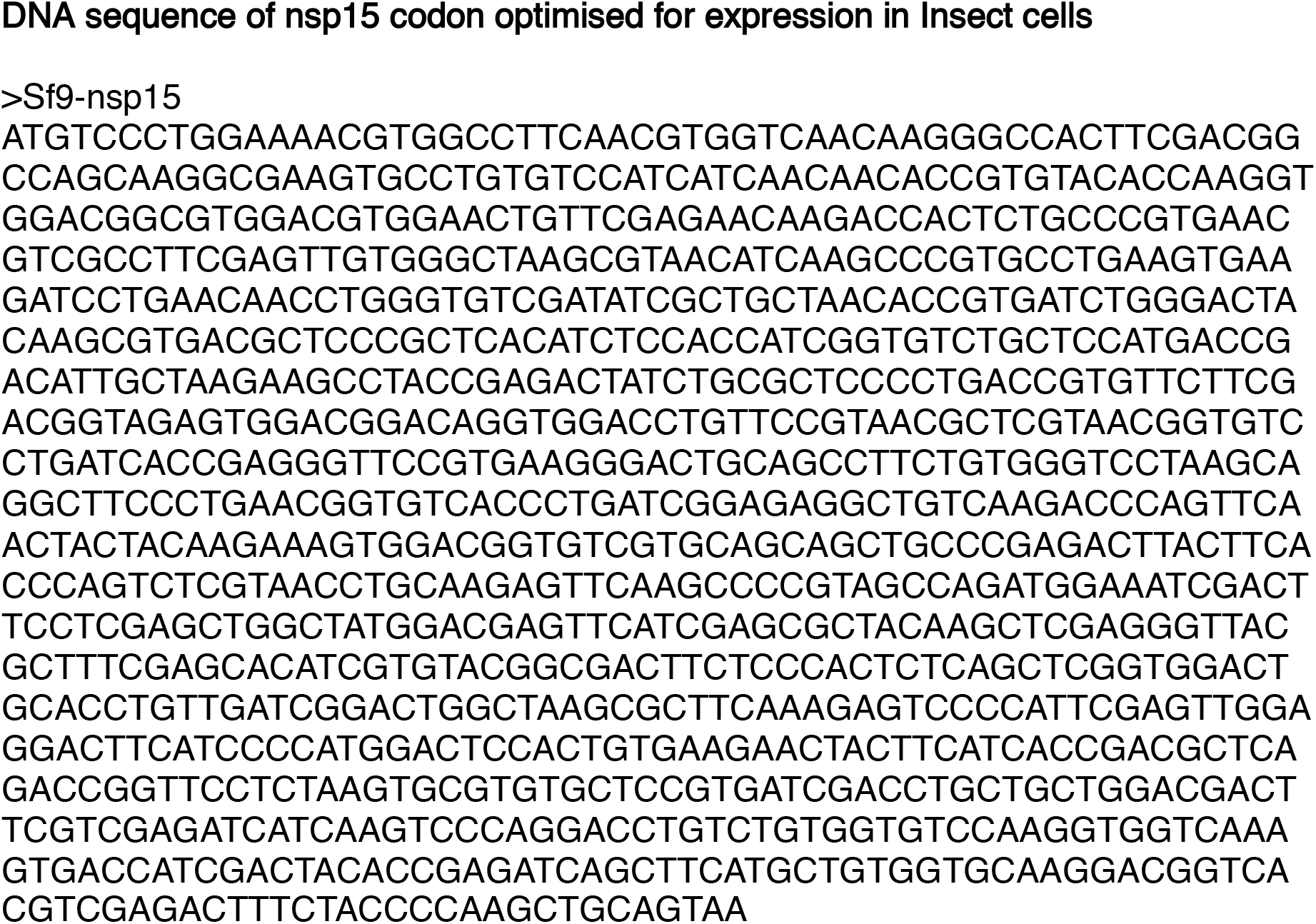
Screen results for known and predicted nsp15 inhibitors.

